# Recent breeding experience improves egg ejection behaviour

**DOI:** 10.1101/2025.01.06.631447

**Authors:** Lisandrina Mari, Anna E. Hughes, Jolyon Troscianko, Oldřich Tomášek, Tomáš Albrecht, Václav Jelínek, Michal Šulc

## Abstract

Recognizing one’s own eggs is crucial for birds, especially for hosts of brood parasites that must identify and reject different-looking parasitic eggs. While birds seem to possess a ‘template image’ of their eggs, whether it is innate or learnt remains unclear. We addressed this question by experimentally inserting either artificial mimetic eggs (ME) or non-mimetic eggs (NME) into the nests of barn swallows (*Hirundo rustica*) during pre-laying stage. Our individually marked population allowed a unique comparison between naïve first-time breeders and experienced females, as well as females with ‘old’ (from a previous season) and ‘recent’ (from the previous breeding attempt within the same season) experience allowing to investigate the role of memory. We found that both naïve and experienced females ejected ME and NME at similar rates, indicating that the template image is neither innate nor learnt and does not play a primary role in egg recognition. Instead, our findings suggest that awareness of own egg-laying is the crucial mechanism at play, and may facilitate nest sanitation behaviour. Finally, we provide the first evidence that this mechanism improves with recent breeding experience. Future studies should investigate the role of non-visual factors that may modulate nest sanitation, such as endocrine regulation.

## Introduction

Whether birds can recognise their own eggs has interested evolutionary biologists for over a century [1,2]. Research has shown that many species benefit from recognizing their eggs appearance. Ground-nesting species may use this knowledge to improve camouflage by selecting nesting surfaces that visually match their eggs [3], while colonial nesters like common murres (*Uria aalge*) are thought to rely on egg appearance to identify their clutches within dense colonies [4,5]. Brood parasites, who exploit host parental care by laying eggs in their nests, strategically choose hosts whose eggs closely resemble their own to avoid detection and rejection by hosts [6,7]. In response, hosts have evolved the capacity to recognize and reject foreign eggs using a “template image” of their own eggs [8]. This mental reference allows hosts to detect parasitic eggs even in the absence of their own [9,10].

Considerable scientific effort has been devoted to understanding whether birds possess an innate template or learn it through breeding experience. Older studies observed anecdotal cases of females rejecting their own eggs after researchers replaced them with non-mimetic experimental eggs, suggesting that birds imprinted on the experimental eggs and later rejected their own [2,11]. A more recent study [12] was unable to resolve this question, as house sparrows (*Passer domesticus*) exhibited almost no rejection behaviour towards experimental eggs, the authors therefore suggested that sparrows might acquire a recognition template during each new breeding attempt.

A key approach to distinguishing whether the template is innate or learnt is to compare naïve females (first-time breeders with no prior experience with their own eggs) with experienced females. While younger females are expected to accept more foreign eggs than older experienced females (in the case the template is learnt), evidence for age-related effects is inconsistent [13–18]. However, this may simply reflect a lack of studies involving truly naïve individuals. A recent study on captive Japanese quails (*Coturnix japonica*) suggested that females learn the appearance of their eggs, as only experienced females were able to lay them on visually similar surfaces [3]. However, authors ambiguously considered females that had previously laid a number of eggs as naïve. Additionally, it has been hypothesized that the egg recognition template, whether innate or learnt, may not be fixed and could be refined during initial or later breeding attempts, though more evidence is needed [20].

This study aims to determine whether birds possess an innate template image of their eggs or acquire it through breeding experience. The barn swallow (*Hirundo rustica*, hereafter swallow) serves as an ideal model species due to its ability to eject model eggs [20], and the unique opportunity to distinguish between naïve first-time breeders (who have never seen their own eggs) and experienced females. This distinction is made possible by the species’ exceptionally high breeding site fidelity [21,22] and our ability to capture and mark all breeding individuals within the study area. To test truly naïve individuals that have never encountered their own eggs, we placed either mimetic egg (ME) or non-mimetic egg (NME) in swallow nests during the pre-laying stage. If the template is innate, even naïve females should eject NME more frequently than ME. If the template is not innate but learnt, we predicted that 1) only experienced females would reject NME more frequently than ME, and 2) experienced females would eject NME more often than naïve females, while ME would be ejected at similar rates by both groups.

As barn swallows usually breed twice a year in our study population and display high breeding-site fidelity, we also explored how the timing of breeding experience – whether from a previous season (‘old’ experience) or within the same season (‘recent’ experience) – affects the ejection of experimental eggs. Since prolonged intervals could lead to memory loss [23,24], we predicted that recent experience with one’s own eggs would increase ejection behaviour more than older experience.

Alternatively, if both naïve and experienced swallows eject ME and NME at similar rates, it would indicate that they do not rely on a template image for egg recognition. Instead, egg recognition could be driven by other mechanisms, such as an awareness of their own egg-laying [25,26]. This ‘onset-of-laying’ mechanism, in which birds reject eggs introduced into the nest before laying their first egg, has been documented in multiple species, including barn swallows [20,27,28]. Our experimental design provides a comprehensive test of these competing hypotheses and offers valuable insights into the mechanisms underlying egg recognition and ejection behaviour.

## Materials and methods

### Study area and general approach

Barn swallows were studied during four breeding seasons from 2021 to 2024 in four farms in the villages Stará Hlína (49° 02′ 21.4″N, 14° 49′ 06.8″E), Břilice (49° 01′ 14.4″N, 14° 44′ 15.3″E), Lužnice (49°3’25.3”N, 14°46’11.4”E) and Lomnice nad Lužnicí (49°4’7.7”N, 14°42’36.7”E) in southern Bohemia, Czech Republic, where they nest inside cattle barns on walls, beams, hanging lamps, or in crevices near ceilings. They arrive in late March, and females begin laying eggs in late April or early May. This population has been monitored since 2010, with all adults and nestlings ringed every year.

During four annual mist-netting sessions (from May to July), previously ringed adults were recaptured, and unringed adults were given unique combinations of one aluminium and three colour rings. All adults were sexed using tail length, brood patch, and cloacal protuberance, weighed, measured, photographed, and sampled for feathers and blood. Swallows in our population exhibit extremely high breeding-site fidelity which is consistent with findings from other populations [21,22]. Therefore, following previous research [29,30], we classified unringed birds as first-year breeders that likely fledged outside our study area during the previous breeding season. Furthermore, detailed nest monitoring throughout the entire season enabled us to distinguish between first and subsequent breeding attempts of individual females. Breeding pairs were photographed at their nests allowing individual identification based on unique ring combinations. Selected breeding attempts were continuously video-recorded with custom Mini CCTV cameras and a digital video-recorder (DVR 4616A ELN AHD lite, Shenzhen DIGIT Co. Ltd, Shenzhen, China).

### Experimental parasitism

To compare ejection rates between naïve and experienced females, we adapted our previous experimental design [20], introducing either a mimetic (ME) or non-mimetic (NME) artificial egg into swallow nests during the pre-laying stage. This stage is ideal for studying the acquisition and use of the template image for egg recognition, as it captures responses before birds see their own eggs, offering valuable insights into whether template image is innate or learnt. During the pre-laying stage, birds may reject any egg present as a part of the ‘onset-of-laying’ mechanism [25–27,31]. However, since we found that swallows often tolerated model eggs inserted at this stage [20], their ejection behaviour may also depend on a template image.

ME closely resembled average swallow eggs in size, weight, and spotting pattern, with colour adjusted for avian perception. NME matched ME in size, weight, and shape but were painted matte blue (Pantone 7459C). Colour (ΔS) and luminance (ΔL) differences from the real model egg were measured using ImageJ [32] with the MICA Toolbox [33]. We consider ME to be excellent mimics of swallow eggs due to their small ΔS and ΔL differences from real eggs (**Figure 1**). This assessment is further supported by our prior research using the same experimental eggs, where swallows accepted 96% of ME, compared to 70% acceptance for NME during the laying stage [20]. Detailed protocols for crafting the experimental eggs and calculation of colour and luminance differences are available in [20].

**Fig 1.**
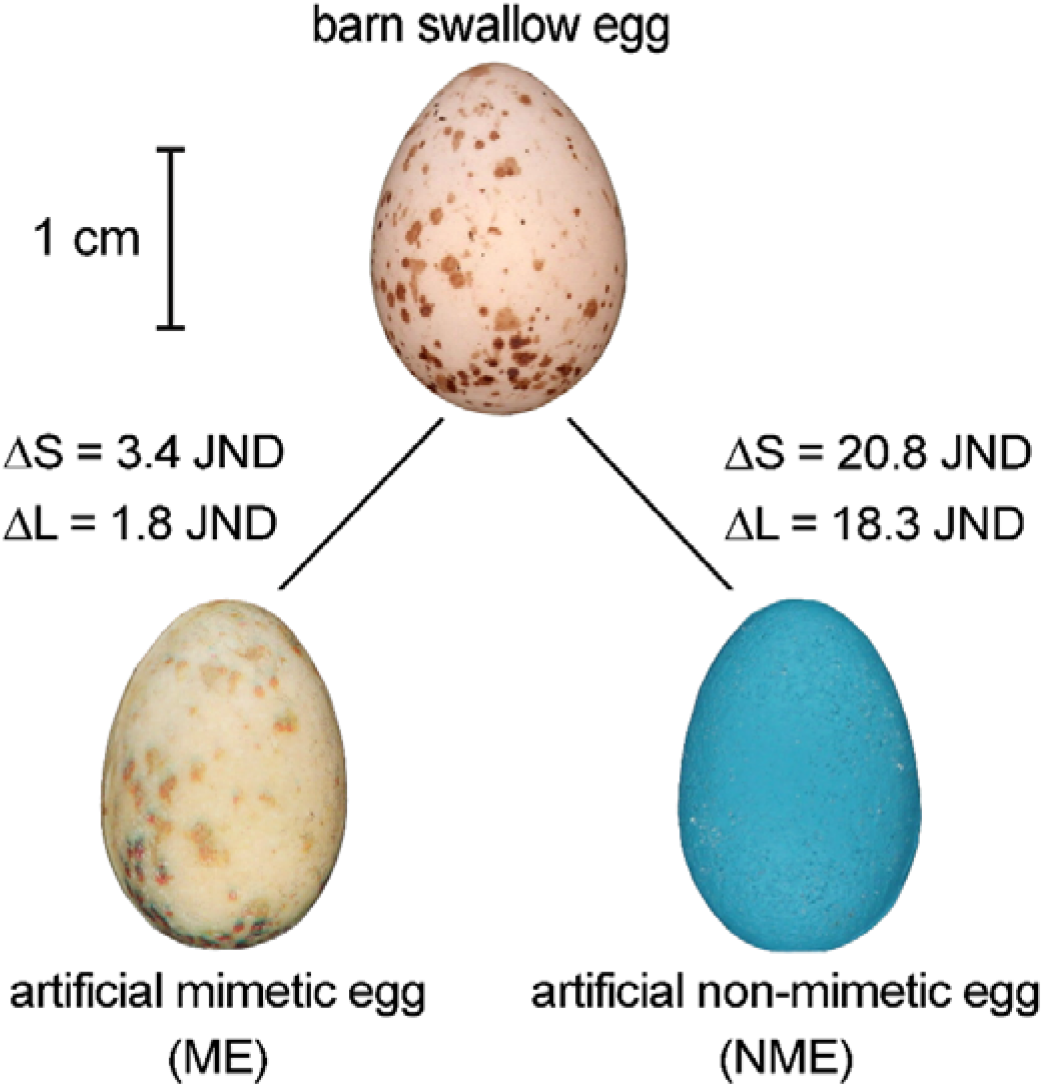
Artificial eggs used in experiments, and a real barn swallow egg that served as a model for creating artificial eggs. Chromatic (ΔS) and luminance (ΔL) differences between the real egg and both ME and NME are in JND values (just noticeable differences) with higher values indicating larger differences. Differences < 3 JNDs indicate colours that are very difficult to distinguish under suboptimal lighting conditions [34–36], which is often the case of swallow nests [20].

Active nests in the pre-laying stage were identified by the appearance of feathers in the nest cup which females use as the final lining material. Then we started our experiment by inserting either ME or NME in the nest. After that, nests were checked daily, with experiments ending when the experimental egg was either ejected or the first swallow egg was laid, in which case the experimental egg was removed and classified as accepted. Presentation durations (time between egg insertion and first-egg laying) averaged 3.6 ± 1.7 (SD) days across experiments (N = 187) and were similar for ME and NME (probability of difference > 0 = 0.435). Ejections occurred within 22.7 ± 17.9 hours (N = 50) on average, with no significant difference between egg types (probability of difference > 0 = 0.358). To ensure consistency, eggs were considered accepted only if they were inserted in the nest at least two days before the first egg was laid. To confirm that ejections were performed by breeding females, we included only those occurring within five days before egg-laying. Video recordings from 14 nests showed that males occasionally participated in ejections (2 cases). However, males never lined nests with feathers and only rarely visited and interacted with nest contents (based on 10 recorded nests). Thus, we assumed most ejections were performed by females and excluded the two male-involved cases from analyses. Our final dataset consists of 187 experiments during which we added either a mimetic (N=83) or a non-mimetic blue egg (N=104) to a swallow nest.

### Statistical analyses

To test our hypotheses, we built two Bayesian hierarchical models. The first model compared the egg ejection response of naïve vs experienced females to determine whether the template image of their own eggs is innate or learnt. Naïve females were defined as first year breeders in their first breeding attempt of the season. These females were considered naïve because they had never laid eggs and likely had no prior experience with swallow eggs in their lifetime. In contrast, experienced females included individuals that had bred at least once at our study site. This group comprised both older females that had bred in previous seasons and first year breeders during their second breeding attempt of the season.

The model analysed the ejection response, coded as a binary outcome (0 = acceptance, 1 = ejection) and modelled using a Bernoulli distribution. Female experience (naïve or experienced) and experimental egg type (ME or NME) were included as categorical predictors, together with their interaction to capture potential combined effects. To account for repeatedly tested females (N = 38), their identity was included as a random factor. We used a normal(0,10) prior for life experience, egg type and the interaction, and a Cauchy(0,2) prior for the random effects.

In the second model, we investigated whether the swallows’ ejection behaviour differed based on whether their experience with own eggs was *recent* or *old. Recent experience* referred to females that had seen their eggs during a previous breeding attempt in the same season, typically about one month earlier (mean ± SD: 33.5 ± 4.6 days; N = 21). *Old experience* corresponded to females that had bred in the previous season, approximately 10-11 months earlier (292.9 ± 19.0 days; N = 26). We thus categorized females experience into three groups: naïve females, females with recent experience and females with old experience. We constructed a full model that included the type of experience, the type of experimental egg (ME or NME), and their interaction as explanatory variables. The random factor (female identity) and priors were set to be the same as in the previous model.

In each model, we tested our specific hypotheses using selected contrasts (e.g. naïve vs. experienced females, ME vs. NME). For each model, we first extracted 1000 draws from the transformed (i.e. on the 0-1 probability scale and not the log odds scale) posterior distributions of conditional means for every possible combination of explanatory factors (which varied depending on the model, see above). We then calculated relevant contrasts by subtracting draws from each other. From these, we calculated the probability of the difference being greater than zero i.e. showing whether the conditions in the contrast of interest reliably show differences in their ejection rates according to the model. Our definition of strong evidence is where the probability of difference is higher than zero with 95% probability (i.e., > 0.95). Due to the lower power of binomial modelling, we further consider a difference higher than zero with 90% probability as non-negligible but weak evidence. All statistical analyses were performed in R 4.4.0 [37] using the *brms* package (2.21.0) [38].

## Results

We found no evidence supporting an innate template image, as naïve females ejected ME and NME at similar rates (probability of difference > 0 = 0.551). Although **Figure 2A** suggests that experienced females showed an overall increase in ejection behaviour compared to naïve females, our model provided only weak evidence for higher ejection rate in experienced females (probability of difference between experienced and non-experienced females > 0 = 0.906). This increase was especially driven by increased ejection of NME rather than ME (probability > 0 = 0.900 for NME vs. 0.762 for ME). Moreover, experienced females did not develop a template image of their eggs, as they ejected NME and ME at similar rates (probability of difference > 0 = 0.699).

**Fig 2.**
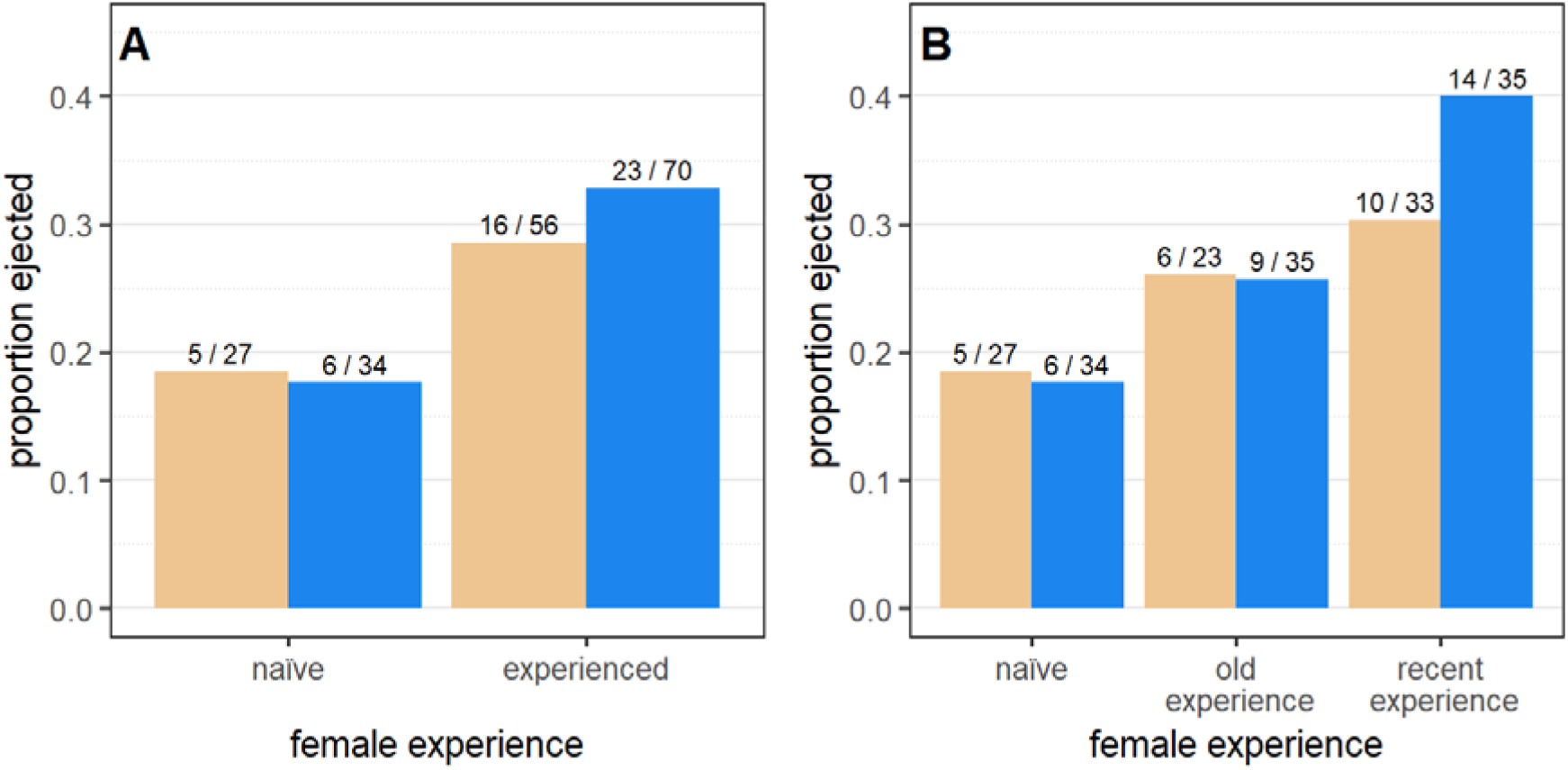
Proportion of experimental eggs (ME or NME) ejected, with A) difference between naïve and experienced females in general, and B) detailed differences among females with old, recent, or no experience (naïve females).

Interestingly, **Figure 2B** indicates that females with recent experience were the most likely to eject model eggs, followed by females with old experience, and finally naïve females. However, we found no strong evidence for a difference in ejection rates between naïve females and females with old experience (probability of difference > 0 = 0.706), or between females with old and recent experience (probability of difference > 0 = 0.858). The only robust evidence was that females with recent experience ejected model eggs more often than naïve females (probability of difference > 0 = 0.956), driven primarily by NME ejection (probability > 0 = 0.96 for NME vs. 0.758 for ME). Similarly to experienced females, females with recent experience did not acquire a template image as they ejected NME and ME at comparable rates (probability of difference > 0 = 0.8).

## Discussion

Our findings demonstrate that barn swallows lack an innate template image of their eggs, as naïve females ejected both ME and NME at similar rates. While a previous study on house sparrows (*Passer domesticus*) suggested the same conclusion [12], researchers experimentally altered the appearance of eggs shortly after laying, leaving the possibility that sparrows may have already seen and imprinted on their own eggs before the manipulation. Furthermore, sparrows showed no egg ejection during the experimental treatments [12], raising concerns on whether the egg manipulation was adequately designed to trigger a recognition.

Although experienced females exhibited increased ejection of both ME and NME compared to naïve females, the increase was only weakly significant for NME. Contrary to predictions, experienced swallows did not eject NME more frequently than ME, indicating they did not acquire a template image through breeding experience either. This result corresponds to our previous study, which suggested that swallows lack a colour template image and instead prioritize shape traits when identifying foreign objects in their nests [20]. In contrast, a similarly designed experiment showed that yellow-bellied prinias (*Prinia flaviventris*) use a template image, though the role of individual experience was not investigated [27]. Additional indirect evidence for a template image comes from studies showing older females are more likely to eject parasitic eggs [13–18]. Although these studies contradict our results, we propose that the absence of a template in our barn swallow population may result from weak brood parasitism pressure. While barn swallows were historically considered conspecific brood parasites [31,39], recent research demonstrates this behaviour is in fact rare [40], and interspecific parasitism is absent in Europe [41]. Consequently, the selection pressure for swallows to develop a template image of their eggs is minimal compared to major hosts of brood parasites (for which the existence of a template image has been suggested [13–18,27]). Additionally, colony-living likely did not drive the evolution of egg recognition in swallows either, unlike other species [4,5]. This may stem from the reliance of swallows on information other than egg phenotype, e.g. spatial memory, to locate their nest, reducing the need for clutch recognition.

Our observation that experienced females increased ejection of both ME and NME suggests that egg recognition relies on other mechanisms than a colour template, such as the awareness of their own egg-laying, also called onset-of-laying mechanism. Studies on barn swallows and cliff swallows (*Petrochelidon pyrrhonota*) show that they eject even mimetic eggs if introduced before the first egg is laid [20,31,42]. Similarly, some brood parasites hosts are more likely to reject parasitic eggs laid in empty nests [27,28, but see 43 for more complex results]. Given the low parasitism level in our population, we propose that the pre-laying ejection behaviour is not a response to parasitism but rather a strategy linked to the frequent reuse of old nests, which may contain unhatched eggs from previous breeding attempts. This aligns with the highly developed nest sanitation behaviour observed in barn swallows [20,44], important for maintaining optimal breeding conditions.

Interestingly, our observations showed that ejection behaviour for both ME and NME was highest in females with recent breeding experience, followed by females with older experience, and finally naïve females. If the onset-of-laying is the primary mechanism underlying egg ejection, this provides the first evidence that awareness of egg-laying improves with breeding experience, particularly recent experience. The improved ejection behaviour in females with the recent breeding experience indicates that attention to nest content may increase as the breeding season progresses. Since the transition between pre-breeding and breeding states involves physiological changes [45], the hormonal makeup of individual females may directly influence ejection decisions, as shown in brood parasite hosts [46–49]. Notably, catecholamines and steroid hormones have been associated with the emergence of sexual behaviours and are known to regulate sensory and cognitive processes [50,51]. As such, females at the onset of their first breeding attempt might be less attuned to the need to sanitize their nests, whereas females in their second breeding attempt have already experienced the hormonal fluctuations that enhance their cognitive abilities. This could also explain why naïve females and females with old breeding experience showed similar ejection rates, as both undergo similar physiological transitions between seasonal states. In addition, increased ejection behaviour in females with recent breeding experience may also reflect a stronger need to remove unhatched eggs, which are more common in reused nests than initial nests of the season (personal observation).

In conclusion, our findings demonstrate that barn swallows lack innate knowledge of their eggs’ appearance and do not acquire it through breeding experience either. This could be attributed to the lack of selective pressures required to drive the evolution of this behaviour. Instead, our results highlight the critical role of the onset-of-laying mechanism, where swallows rely on the awareness of their egg-laying to maintain nest sanitation [20,44]. This mechanism improves with breeding experience and reaches its peak during second breeding attempts, during which removing unhatched eggs from previous nesting attempts may be particularly important. Future research should investigate the interplay between endocrine regulation and cognition, and integrate hormonal and behavioural data to establish links between seasonality, physiology and nest sanitation. This approach could reveal overlooked mechanisms that improve egg ejection behaviour, the main defence against brood parasitism.

## Acknowledgements

We thank our students and colleagues, including K. Vlčková, M. Frýbová, H. Hal’amová, M. Janča, Š. Kadlecová, K. Kopecká, P. Kopková, M. Kuba, K. Míčková, K. Pačandová, L. Pazdera, G. Večeřová,

J. Týblová, B. Počtová Vodičková, J. Záleská, H. Zdobinská and L. Zemanová for their invaluable assistance with fieldwork. We are also grateful to the managers of the cattle farms that kindly allowed us to conduct our fieldwork on their grounds.

## Author contributions

MŠ, LM, AEH and VJ conceived the ideas; MŠ, LM, AEH, JT and VJ designed methodology; MŠ, VJ, OT and TA collected data; AEH, LM, MŠ and OT performed statistical analyses; MŠ and LM wrote the initial draft of the manuscript. All authors contributed to the drafts and gave final approval for publication.

## Ethical approval

We declare that all experiments performed for this study were approved by the animal and ethics representatives of The Czech Academy of Sciences and nature conservation authorities (62065/2017-MZE-17214 and MZP/2020/630/964). The fieldwork adhered to the Czech Law on the Protection of Animals against Mistreatment (licence no. CZ03971 and CZ04122).

